# Pathogenic mutations in LRRK2 sequester Rab8a to damaged lysosomes and regulate transferrin-mediated iron uptake in microglia

**DOI:** 10.1101/2020.07.27.219501

**Authors:** Adamantios Mamais, Natalie Landeck, Rebekah G. Langston, Luis Bonet-Ponce, Nathan Smith, Alexandra Beilina, Alice Kaganovich, Manik C. Ghosh, Laura Pellegrini, Jillian H. Kluss, Ravindran Kumaran, Ioannis Papazoglou, Nunziata Maio, Changyoun Kim, David C. Gershlick, Mark R. Cookson

## Abstract

Mutations in leucine-rich repeat kinase 2 (LRRK2) cause autosomal dominant Parkinson’s disease (PD) while polymorphic LRRK2 variants are associated with sporadic PD. PD-linked mutations increase LRRK2 kinase activity and induce neurotoxicity *in vitro* and *in vivo*. The small GTPase Rab8a is a LRRK2 kinase substrate and is involved in receptor-mediated recycling and endocytic trafficking of transferrin, but the effect of PD-linked LRRK2 mutations on the function of Rab8a are poorly understood. Here, we show that gain-of-function mutations in LRRK2 induce sequestration of endogenous Rab8a into lysosomes in cells while pharmacological inhibition of LRRK2 kinase activity reverses this phenotype. Furthermore, we show that LRRK2 mutations drive accumulation of endocytosed transferrin into Rab8a-positive lysosomes leading to a dysregulation of iron transport. LRRK2 has been nominated as an integral part of cellular responses downstream of proinflammatory signals and is activated in microglia in post-mortem PD tissue. Here, we show that iPSC-derived microglia from patients carrying the most common LRRK2 mutation, G2019S, mistraffic transferrin to lysosomes proximal to the nucleus in proinflammatory conditions. Furthermore, G2019S knock-in mice show significant increase in iron deposition in microglia following intrastriatal LPS injection compared to wild type mice, accompanied by striatal accumulation of ferritin. Our data support a role of LRRK2 in modulating iron uptake and storage in response to proinflammatory stimuli in microglia.

## Introduction

Missense mutations on the LRRK2 gene cause autosomal dominant Parkinson’s disease (PD), while common genetic variants identified by genome-wide association studies have been linked to idiopathic PD and inflammatory diseases (1–3). LRRK2 is a multidomain enzyme and PD-linked mutations are localized to its enzymatic domains, namely the kinase domain and Ras of complex proteins and C-terminal of ROC (ROC-COR) domains. LRRK2 has been linked to a number of cellular pathways including autophagy, lysosomal processing, inflammation and vesicular trafficking (4). LRRK2 is expressed in immune cells and reports suggest a role in microglial activation and an effect of PD-linked mutations on cytokine release and inflammation (5–7). The kinase activity of LRRK2 is thought to drive pathology in disease, and thus is being targeted pharmacologically as a possible therapy (4).

A specific role of LRRK2 on vesicular trafficking has been nominated in recent years and a number of studies have highlighted how downstream signaling pathways are affected by PD-linked mutations. LRRK2 can phosphorylate a subset of Rab GTPases, including Rab8, Rab10 and Rab29, at a conserved motif and this phosphorylation can regulate Rab activity and association with effector molecules (8,9). Rab GTPases control the spatiotemporal regulation of vesicle traffic, and are involved in vesicle sorting, motility and fusion (10). Different Rab GTPases exhibit selectivity for different cellular compartments thus conferring membrane identity that governs secretory and endocytic pathways (11). Our lab and others have shown that LRRK2 associates with Rab29, a Rab GTPase that is involved in vesicular transport and protein sorting (12,13). Rab29 is a candidate risk gene for sporadic PD (14) and can mediate recruitment of LRRK2 to the trans-Golgi network, inducing LRRK2 activity and membrane association (12,15,16). LRRK2 can phosphorylate Rab8 and Rab10, and reports suggest downstream effects on lysosomal integrity. We, and others, have reported phosphorylation-dependent recruitment of Rab10 onto damaged lysosomes by LRRK2, in a process that orchestrates lysosomal homeostasis (17,18).

Rab8a is involved in a number of cellular functions including cell morphogenesis, neuronal differentiation, ciliogenesis and membrane trafficking (19–21). Rab8a regulates recycling of internalized receptors and membrane trafficking to the endocytic recycling compartment (ERC) (20,22). Rab8a has been shown to interact directly with the transferrin receptor (TfR) and regulate polarized TfR recycling to cell protrusions (23) and transfer between cells through tunneling nanotubes (24). Furthermore, Rab8a-depleted cells fail to deliver internalized TfR to the ERC (23). Rab8a activity is controlled by its association with Rabin8, a guanine exchange factor (25). Phosphorylation of Rab8a by LRRK2 inhibits its association with Rabin8 and thereby modulates Rab8a activity (8). The G2019S LRRK2 mutation enhances Rab8a phosphorylation, mediating defects on EGFR recycling and endolysosomal transport (26).

Although these data indicate that Rab8a regulates exocytic and recycling membrane trafficking at the ERC, how phosphorylation of Rab8a affects downstream biology in PD-relevant cell types remains unclear. Here we identify a novel role of LRRK2 in mediating Rab8a-dependent TfR recycling and show *in vivo* effects of LRRK2 mutation at the endogenous level on iron homeostasis in microglia. Expression of mutant LRRK2 induces sequestration of Rab8a to lysosomes and dysregulates transferrin recycling in a Rab8a-dependent manner. In iPSC-derived microglia carrying the G2019S LRRK2 mutation, transferrin retains association with lysosomes in proinflammatory conditions. Finally, we show that G2019S LRRK2 knock-in mice exhibit increased iron accumulation within microglia in the striatum in response to neuroinflammation compared to WT controls. These data support a role of LRRK2 in modulating iron uptake and storage in response to proinflammatory stimuli in microglia.

## Results

### Pathogenic mutants of LRRK2 sequester Rab8a to damaged lysosomes

In order to study endogenous Rab8a in cells, we first validated commercially available antibodies for western blotting and immunofluorescence using siRNA-mediated knockdown (Fig S1). Using the validated antibody tools we examined the intracellular localization of Rab8a in the context of LRRK2 mutations. Different LRRK2 constructs carrying pathogenic mutations were transiently expressed in Hek293FT cells and endogenous Rab8a was visualized (Fig 1 and Fig S2 for imaging of all mutants). Endogenous Rab8a plays a part in receptor recycling and localizes in tubular recycling endosomes (20). Consistent with such a function, in cells expressing WT LRRK2, Rab8a localized predominantly in tubular membranes not associated with Lamp2-positive lysosomes (Fig 1A). However, expression of R1441C or G2019S LRRK2 variants induced sequestration of endogenous Rab8a into enlarged Lamp2-positive lysosomes (Fig 1A). Other pathogenic mutations, but not the kinase-dead variant K1906M, also induced relocalization of Rab8a to lysosomes (Fig S2). The latter result suggested that the recruitment of Rab8a to lysosomes might be kinase dependent. To test this hypothesis, cells were treated with the LRRK2 kinase inhibitor MLi-2 for one hour prior to fixation and staining. LRRK2 kinase inhibition rescued the phenotype, restoring association of Rab8a with tubular recycling endosomes (Fig 1A). To quantify LRRK2-driven Rab8a recruitment in an unbiased manner, we used a high-content imaging system (Cellomics, Thermo Fisher). The number of sequestered Rab8a structures per cell was counted revealing a convergent phenotype for all PD-linked LRRK2 genetic variants with an increase in Rab8a recruitment that was rescued by MLi-2 (Fig 1B). High-throughput colocalization analysis showed that all pathogenic LRRK2 variants induced association of sequestered Rab8a with LRRK2 (Fig 1C) and Lamp2 (Fig 1D) while the kinase dead LRRK2 mutant did not. Both recruitment of Rab8a by LRRK2 and colocalization with Lamp2 were rescued by MLi-2 treatment (Fig 1C, D). In our experiments, about 50% of enlarged vesicles that were positive for LRRK2 and Rab8a were also positive for Lamp2 in cells expressing mutant LRRK2. To examine whether mutant LRRK2 and Rab8a are recruited to the lysosomal lumen or membrane, cells expressing G2019S LRRK2 were analyzed by super-resolution microscopy (Airyscan), revealing a clear recruitment of both proteins to the membrane of Lamp1-positive lysosomes (Fig 1E).

**Figure 1.**
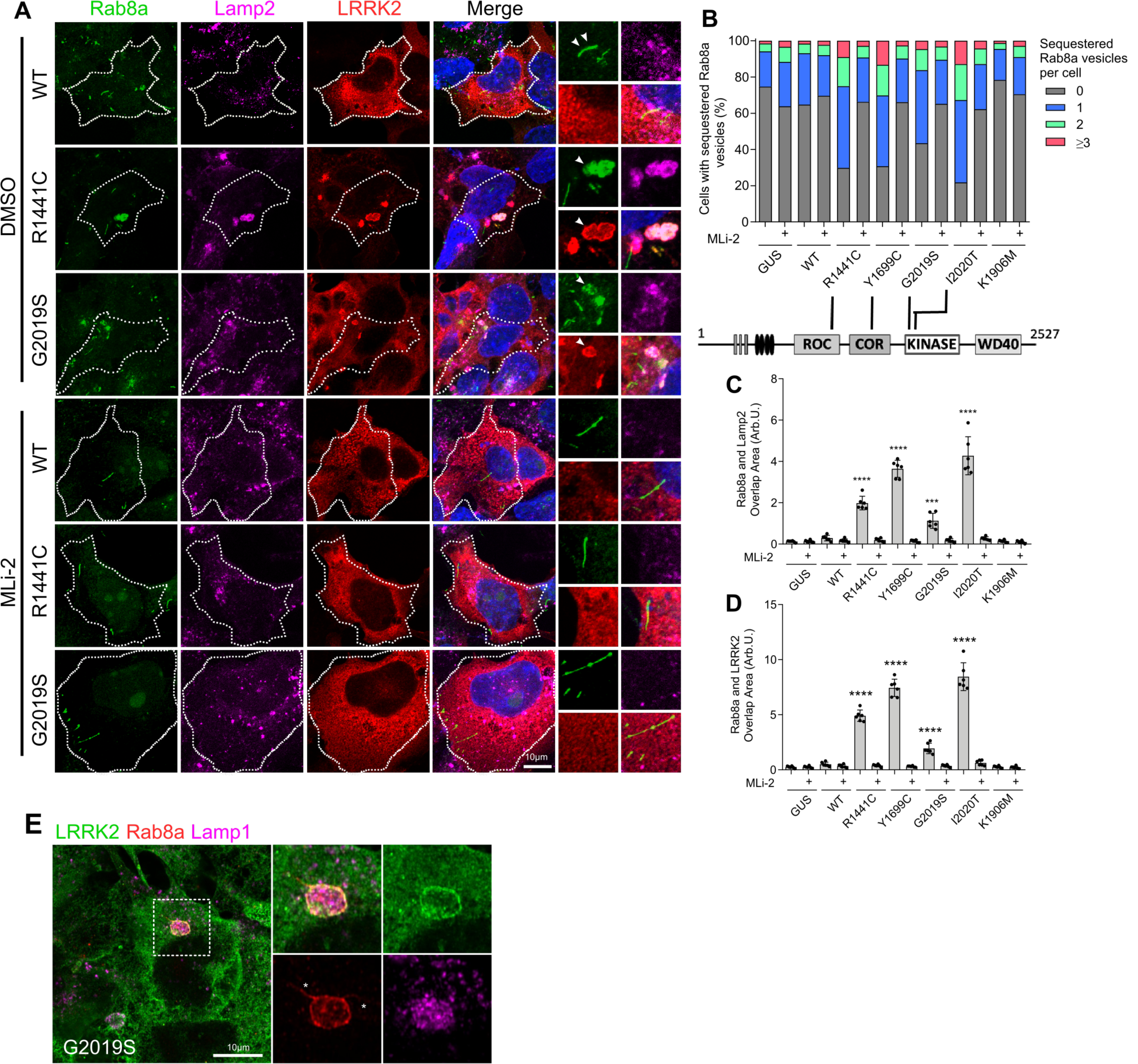
Pathogenic mutations of LRRK2 sequester endogenous Rab8a to lysosomes in a kinase-dependent manner. (A) Endogenous Rab8a is sequestered to Lamp2-positive lysosomes in cells expressing R1441C or G2019S LRRK2, while this is rescued by MLi-2 treatment. (B) Quantitation of Rab8a sequestration by high-content imaging reveals a significantly higher percentage of cells exhibiting sequestered Rab8a (>4µm^2^ foci) in cells expressing R1441C, Y1699C, G2019S and I2020T LRRK2 compared to WT LRRK2, while this is ameliorated by MLi-2. (C, D) Rab8a shows increased colocalization with Lamp2 and LRRK2 in cells expressing LRRK2 PD-linked mutants compared to WT or K1906M LRRK2 and this is rescued by MLi-2 treatment (C; N=6 wells of >800 cells counted per well; two-way ANOVA; treatment: p<0.0001, F (1, 70) = 489.2; genotype: p<0.0001, F (6, 70) = 97.51, D; two-way ANOVA; treatment: p<0.0001, F (1, 70) = 937.7; genotype: p<0.0001; F (6, 70) = 189.8; SD bars are shown). (E) Super-resolution image of endogenous Rab8a and over-expressed G2019S LRRK2 localizing at the lysosomal membrane.

Previous studies have shown that LRRK2 can localize to enlarged lysosomes along with Rab GTPases (18). Furthermore, we have shown that LRRK2 can be recruited to damaged lysosomes that have low degradative capacity (17). In the current model system where we see mutant LRRK2 recruiting Rab8a to Lamp1/Lamp2-positive lysosomes, those LRRK2 structures were found to be Cathepsin D negative (Fig 2A and 2B for Airyscan images). This result suggested that Rab8a was recruited specifically to damaged lysosomes. To further test this hypothesis, we treated cells with the lysosomal destabilizing agent LLOMe that interacts with the lysosomal membrane and luminal hydrolases. Using super-resolution microscopy, we found that LLOMe induced recruitment of WT LRRK2 to the membrane of enlarged lysosomes as previously shown (17) (Fig 2C). Endogenous Rab8a was also recruited to the lysosomal membrane in WT LRRK2 cells after treatment with LLOMe (Fig 2D). Furthermore, the cytosolic scaffolding protein JIP4 was also recruited to the same structure suggesting mistrafficking of factors involved in vesicle-mediated transport (Fig 2E). These data suggest that Rab8a is recruited to damaged lysosomes by mutant LRRK2 in a kinase-dependent manner.

**Figure 2.**
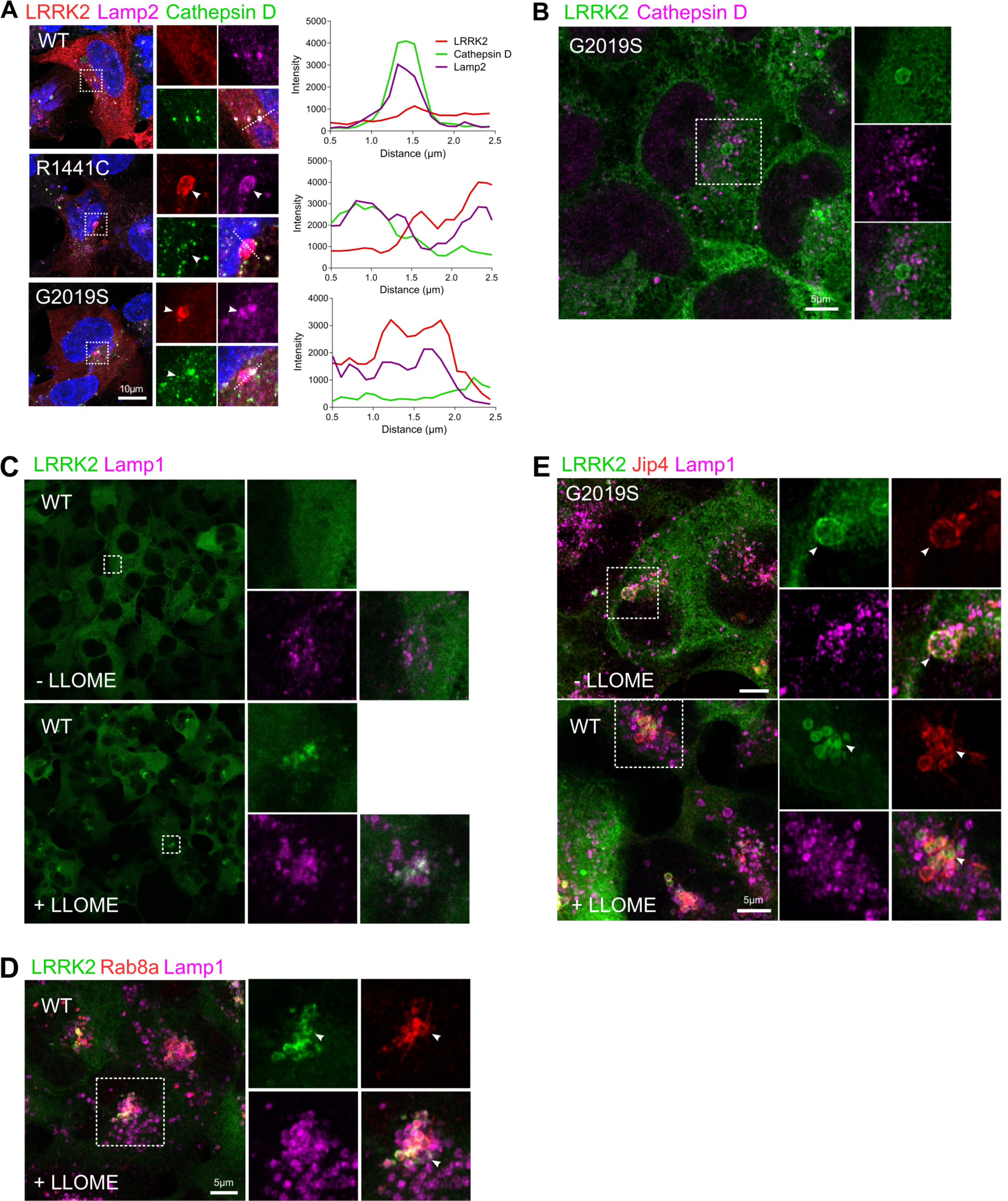
LRRK2 and Rab8a are recruited to the membrane of damaged lysosomes. (A) Transiently-expressed mutant LRRK2 is sequestered to lysosomes that are cathepsin D negative. (B) Super-resolution image of Hek293T cells stably expressing GFP G2019S LRRK2 showing LRRK2 recruitment to the membrane of cathepsin D negative lysosomes. Inducing lysosomal membrane rupture by LLOMe treatment sequesters WT LRRK2 to lysosomes (C) along with endogenous Rab8a (D). LRRK2 G2019S mutation or LLOMe treatment of WT LRRK2 expressing cells induces Jip4 recruitment to lysosomes (E).

### LRRK2-mediated T72 Rab8a phosphorylation blocks interaction with Rabin8 but not MICAL-L1

The above data suggest that Rab8a can move from its normal location at the ERC to the lysosome after expression of LRRK2 mutations. Given that all LRRK2 mutations are proposed to increase Rab phosphorylation, at T72 for Rab8a (8), we reasoned that the mechanism of relocalization might be related to this phosphorylation event. The pT72 Rab8a antibody that was available during this study was not specific to Rab8a and cross-reacted with Rab3A, Rab10, Rab35 and Rab43, which have a high degree of sequence conservation (for datasheet refer to ab230260; Abcam). Therefore, using a tagged version of Rab8a, we found that exogenous expression of R1441C, Y1699C, G2019S or I2020T LRRK2 induced a significant increase in Rab8a phosphorylation at T72 compared to WT or the kinase-dead variant K1906M and this was blocked by MLi-2 treatment (Fig 3A, 3B).

**Figure 3.**
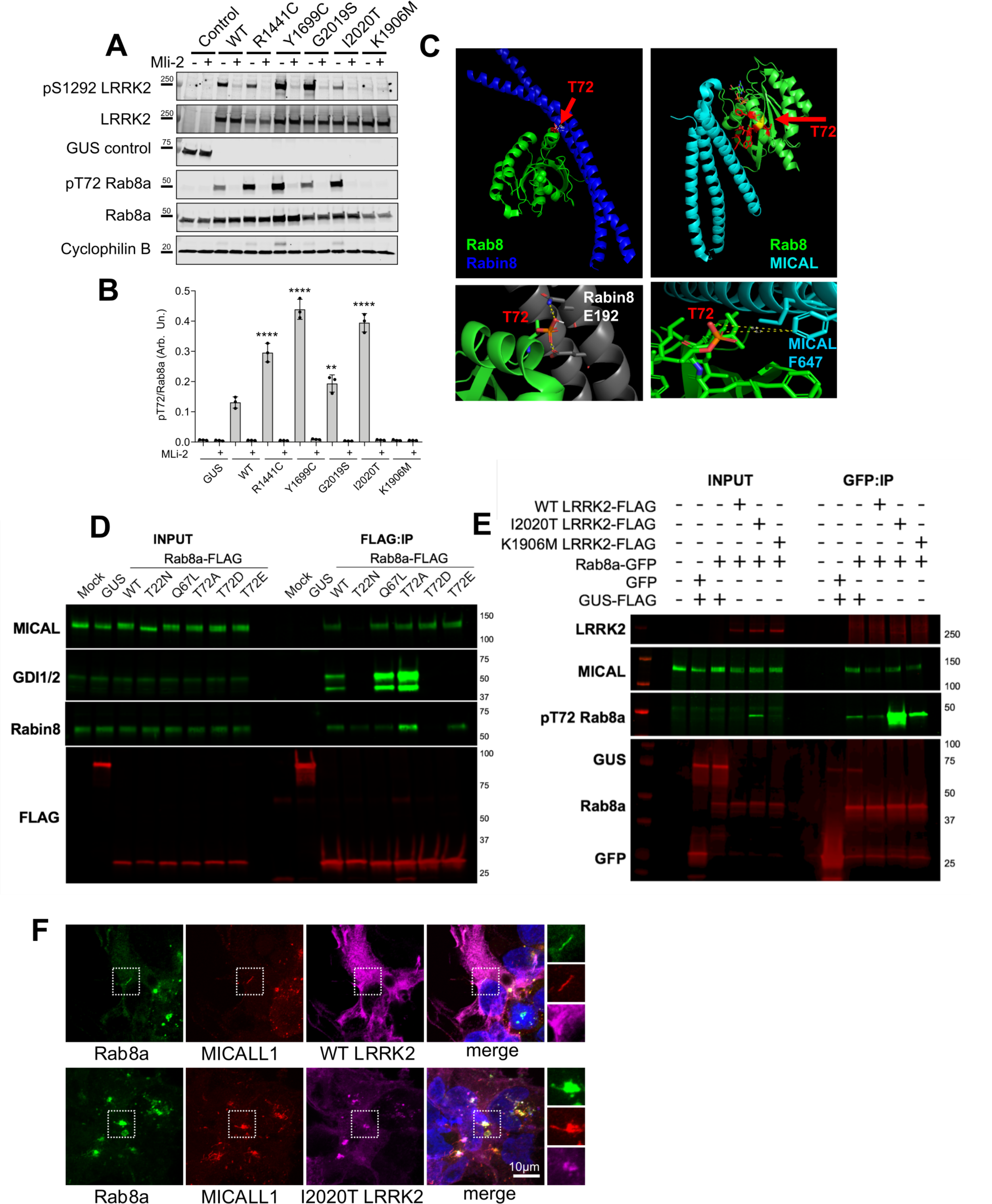
LRRK2-mediated T72 Rab8a phosphorylation blocks interaction with Rabin8 but not MICALL1. (A) T72 Rab8a phosphorylation is significantly increased in cells expressing R1441C, Y1699C, G2019S and I2020T LRRK2 and this is rescued by inhibiting LRRK2 kinase activity (B, two-way ANOVA; N=3; treatment: p<0.0001, F (1, 28) = 1473, genotype: p<0.0001, F (6, 28) = 157.1). (C) Structural modelling predicts that T72 phosphorylation on Rab8a will inhibit the interaction between Rab8a and its GEF Rabin8 (T72 is between 2.8-9.9 A from the closest glutamate on Rabin8). In contrast, T72 phosphorylation lies between 6.7-8.4 A from the closest phenylalanine at MICALL1 suggesting no inhibitory effect. These predictions were validated biochemically by co-IPs (D). Endogenous Rabin8 and GDI1/2 co-immunoprecipitated with exogenously expressed WT Rab8a as well as the phospho-null mutant T72A variant (D). The phospho-mimetic T72D and T72E Rab8a constructs showed decreased interaction with Rabin8 and GDI1/2. MICALL1 associated with WT, phospho-null and phospho-mimetic T72 Rab8a mutants but not with the T22N GTP-binding deficient variant. In a similar setup the LRRK2-driven T72 phosphorylation was investigated (E). T72 Rab8a hyperphosphorylation did not hinder interaction with MICALL1 in cells. (F) Mutant LRRK2 sequesters both Rab8a and interacting partner MICALL1 in cells away from tubular recycling endosomes.

Having established that Rab8a phosphorylation at T72 is increased by all pathogenic LRRK2 mutations, we next considered which effector proteins might be important for functional recruitment from the ERC to lysosomes. We focused on the activating GEF, Rabin8, and MICAL-L1 which localizes in tubular endosomes and is essential for efficient endocytic trafficking from the ERC to the plasma membrane (20,22). Modelling the interaction between Rab8a and Rabin8, we predicted that the Threonine at position 72 on Rab8a resides 2.8-3.9 A away from the neighboring Glutamate (E192) on Rabin8 (Fig 3C). Addition of a phosphate group on T72 will likely hinder the interaction with the negatively charged Glutamate therefore destabilizing Rab8a-Rabin8 interaction. Threonine 72 of Rab8a is closest to a Phenylalanine (F647) on MICAL-L1 at a 6.7-8.4 A distance suggesting that phosphorylation at this site will not affect this interaction. These modeling efforts suggest that Rab8a phosphorylation would be predicted to diminish Rabin8-mediated activation of Rab8a whereas binding to MICAL-L1 should be preserved. To validate these predictions biochemically, we performed co-immunoprecipitation experiments. Endogenous MICAL-L1 co-immunoprecipitated with exogenously expressed WT Rab8a as well as with the GTPase inactive Q67L, phospho-null T72A, phospho-mimetic T72D and T72E Rab8a variants, as predicted (Fig 3D). The T22N Rab8a mutation which renders the protein unable to bind GTP decreased association with MICAL-L1, consistent with the literature (22). The interaction of Rab8a with the guanosine nucleotide dissociation inhibitor GDI1/2 was blocked by the inactive T22N mutant and enhanced in the constitutively active GTP-bound Q67L variant and T72A variants. Rabin8 showed enhanced association with the phospho-null T72A Rab8a variant, consistent with our model. To investigate the effect of LRRK2-dependent Rab8a phosphorylation on interactions, we performed a similar experiment co-expressing Rab8a with WT, kinase-hyperactive I2020T and kinase dead K1906M LRRK2. In these conditions Rab8a was T72 hyperphosphorylated by I2020T LRRK2 and this modification did not hinder association with endogenous MICAL-L1 (Fig 3E). To investigate the retention of this association in the context of Rab8a recruitment to lysosomes, we stained for Rab8a and MICAL-L1 in cells expressing LRRK2. Cells expressing WT LRRK2 showed colocalization of Rab8a with MICAL-L1 in tubular membranes while I2020T LRRK2 expression resulted in the recruitment of both proteins to LRRK2-positive structures consistent with the lysosomal phenotype described above (Fig 3F). Our data suggest that mutant LRRK2 can phosphorylate Rab8a inducing recruitment of both Rab8a and MICAL-L1 away from the ERC to damaged lysosomes.

### Mutant LRRK2 sequesters Rab8a leading to transferrin mistrafficking and accumulation of intracellular iron

Given the above data showing that LRRK2 phosphorylation prevents Rab8a activation and redirects Rab8a and MICAL-L1 away from the ERC, we speculated that these events would then lead to a defect in Rab8a-mediated recycling. To evaluate this hypothesis, we examined the recycling of the transferrin receptor 1 (TfR), which is known to depend on Rab8a function (23). Cells expressing LRRK2 genetic variants were stained for endogenous TfR and analyzed (Fig 4A). TfR localized in distinct cytoplasmic vesicles in cells expressing WT LRRK2, whereas LRRK2 pathogenic mutants showed a clustered TfR localization that associated with the LRRK2 structures we had previously identified as lysosomes. In fact, recruitment of endogenous Rab8a into vesicles positive for TfR and LRRK2 was validated by super-resolution microscopy (Fig S3). The distribution of TfR vesicles in LRRK2-expressing cells was analyzed in Imaris (Bitplane) where vesicles were rendered to spots throughout z-stack 3D reconstructed images and distances between them were measured (Fig 4A). TfR vesicles were significantly less dispersed in cells expressing LRRK2 mutants compared to WT LRRK2 expressing cells (Fig 4B, 4C). This altered distribution prompted us to test whether mutant LRRK2 functionally affected Rab8a-dependent Tf recycling. Cells expressing WT or mutant LRRK2 were preloaded with Alexa Fluor 568-Tf and visualized at T10 minutes of recycling in fresh media. Mutant LRRK2 expressing cells sequestered Tf in enlarged vesicles that were labelled with LRRK2 and Rab8a (Fig 4D). In a time-course experiment, LRRK2 mutant expressing cells plateaued at higher Tf levels compared to WT at T15-20 mins as monitored by high-throughput imaging, pointing to Tf accumulation driven by dysregulated recycling (Fig 4E). Tf that was sequestered by mutant LRRK2 also showed partial co-localization with Lamp2 (Fig 4F), suggesting that the excess Tf accumulates in lysosomes. To test whether Tf accumulation by mutant LRRK2 was Rab8a-dependent, the localization of internalized 568-Tf in I2020T LRRK2 expressing cells was investigated following Rab8a siRNA knock-down. Knock-down of Rab8a expression rescued the increase in the proportion of Tf that co-localized with LRRK2 vesicles relative to total internalized Tf caused by mutant LRRK2 (Fig 4G, 4H). ICP-MS analysis revealed significantly elevated iron levels in cells stably-expressing G2019S LRRK2 compared to WT LRRK2 (Fig 4I). Collectively, these data suggest dysregulation of Tf-mediated iron uptake and iron homeostasis by LRRK2 mutations as a consequence of altered Rab8a localization away from the ERC and towards lysosomes.

**Figure 4.**
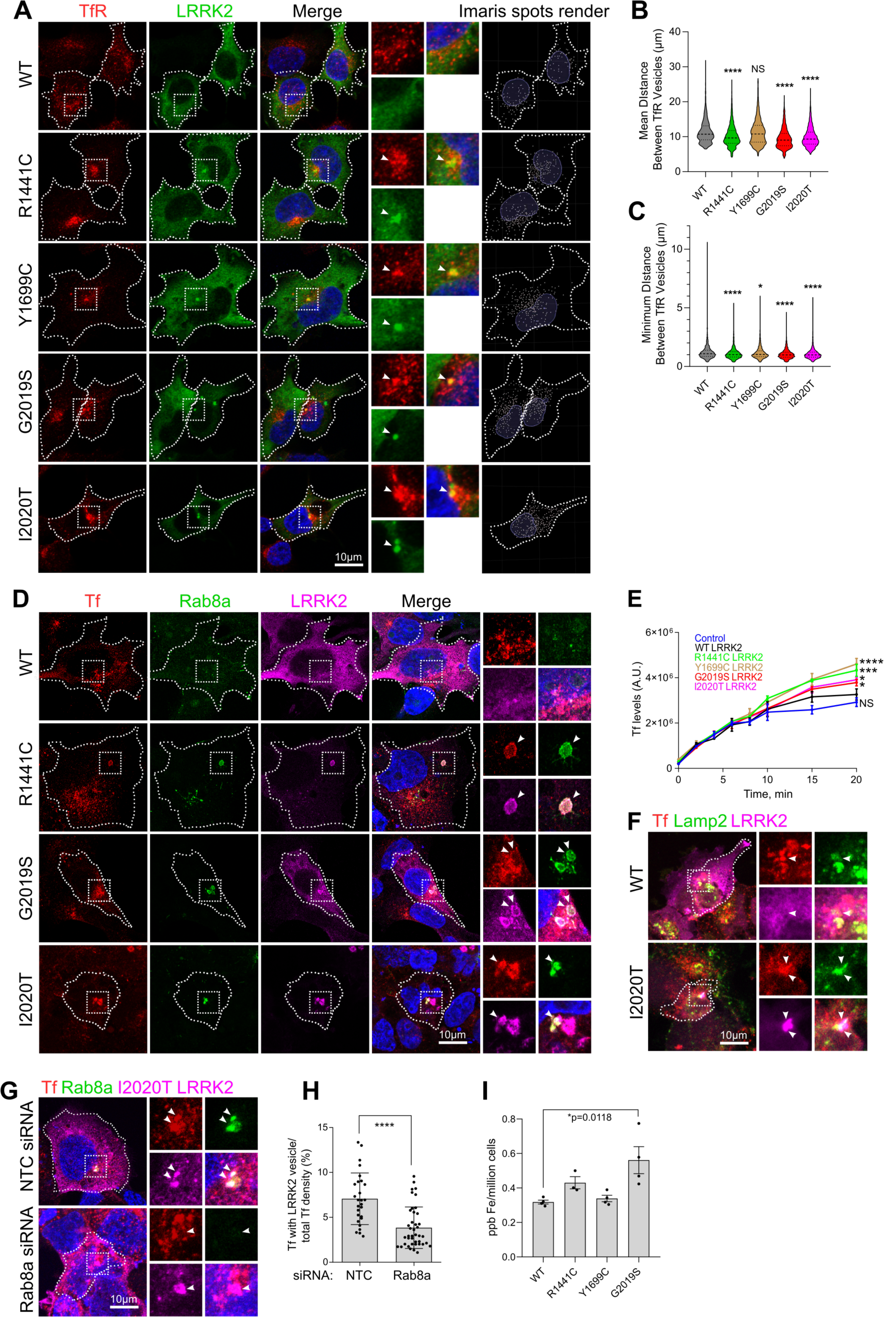
Mutant LRRK2 sequesters TfR and dysregulates transferrin recycling. Cells exogenously expressing WT and mutant LRRK2 constructs were stained for TfR and visualized by super-resolution microscopy. TfR vesicles were analyzed by spot rendering in Imaris and the mean and minimum distances between spots were plotted, revealing that TfR vesicles are clustered closer together in mutant LRRK2 expressing cells compared to WT LRRK2 (B, C). (N>2000 vesicles were counted in at least 20 cells per construct, *P<0.05, ****P<0.0001, one-way anova with Tukey’s post-hoc, B: F(4, 13334) = 46.14, C: F (4, 13336) = 194.1). (D) Hek293FT cells expressing LRRK2 genetic variants were incubated with Alex Fluor 594-conjugated transferrin and visualized. Internalized transferrin was sequestered by mutant LRRK2 along with endogenous Rab8a in large vesicles (D). Time-course of transferrin uptake by high-content imaging revealed accumulation of internalized transferrin in cells expressing LRRK2 mutants compared to WT LRRK2 expressing cells (E: T=20mins, one-way ANOVA, Tukey’s post-hoc, *p<0.05, ***P=0.0005, ****P=<0.0001). Expressing I2020T LRRK2 in cells sequestered Tf to lysosomes (F). Knocking-down Rab8a expression in cells by siRNA rescued the sequestration of transferrin by I2020T LRRK2 exogenous expression (G, H). (N=27 cells for NTC siRNA, N=41 cells for Rab8a siRNA, two-tailed student t-test, Mann Whitney U, ****P<0.0001). Intracellular iron levels were elevated in cells expressing G2019S LRRK2 compared to WT LRRK2 as measured by ICP-MS (I) (N=4, two-way ANOVA with Tukey’s post hoc, *p=0.018, F (3, 11) = 6.225). [SD bars are shown].

### Endolysosomal and iron-binding gene expression in microglia is modulated by inflammation, *in vitro* and *in vivo*

LRRK2 is expressed in microglia and has been linked to inflammation, cytokine release and phagocytosis. Our data presented here show that LRRK2 associates with lysosomes and that LRRK2 mutations dysregulate Tf-dependent iron uptake mechanisms. Glial cells are known to be iron rich and microglia activation state is integrally linked to brain iron content (27). Proinflammatory stimuli are known to induce uptake of extracellular iron by microglia and conversely the iron status of their environment can modulate their activation. Given the role of LRRK2 in inflammation, we asked whether endolysosomal processes and iron homeostasis might converge in proinflammatory conditions. We addressed this hypothesis using two genome-scale approaches. First, we mined our previously described RNA-Seq dataset (28) of primary microglia treated with LPS or preformed α-synuclein fibrils in culture (Fig 5A). Analysis of the shared hits between the two treatments using gene ontology revealed enrichment for endosomal and lysosomal pathways (Fig 5B, 5C). Unsupervised hierarchical clustering of differential gene expression separated the controls from the two treatment groups (Fig 5D). These data demonstrate that genes involved in the endolysosomal system are consistently regulated by inflammatory stimuli.

**Figure 5.**
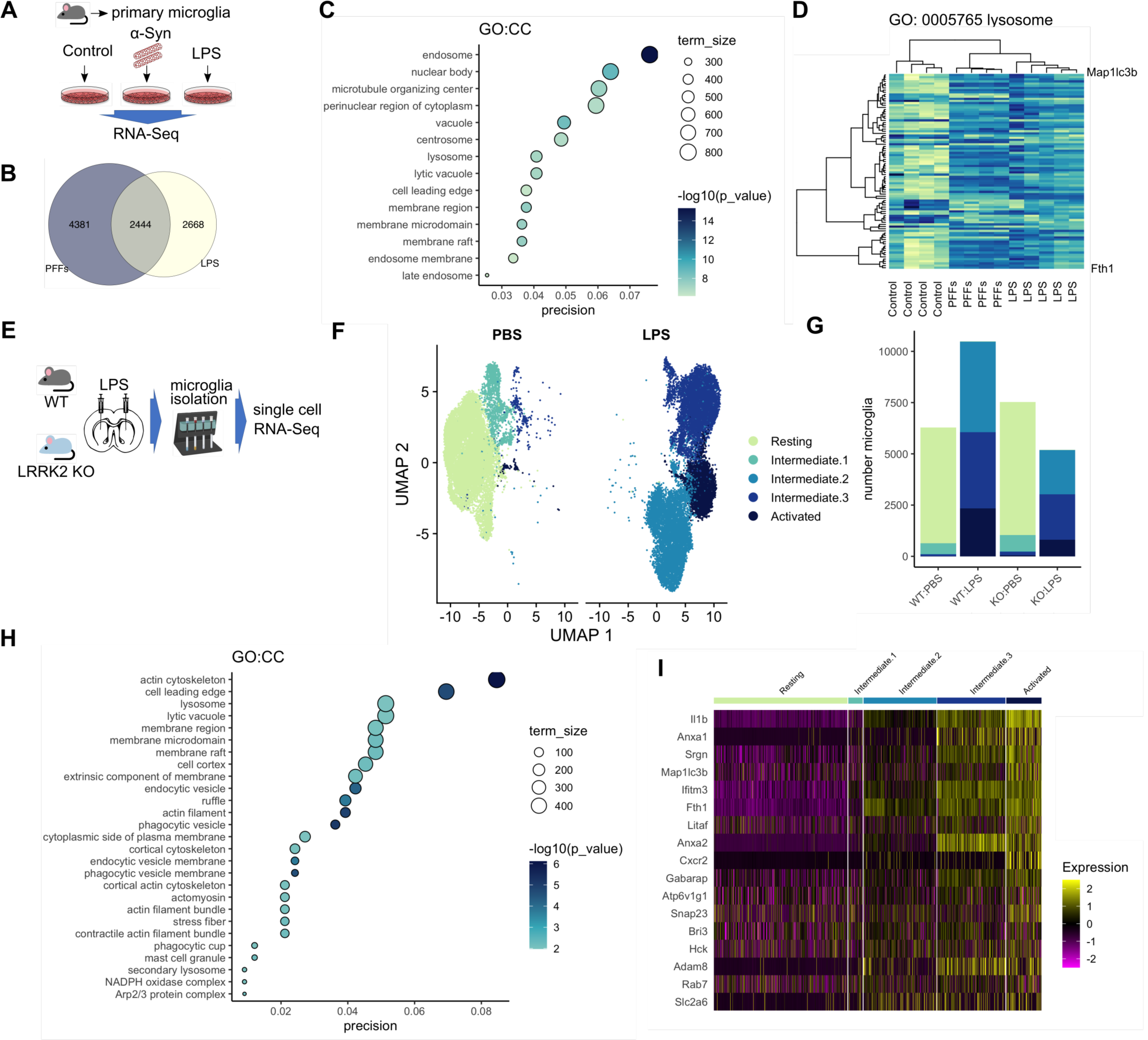
Neuroinflammation remodels endolysosomal gene expression in microglia, *in vitro* and *in vivo*. (A) Outline of RNA Seq experiment: primary mouse microglia cultures were incubated with LPS or α-synuclein fibrils and transcriptomic profiles were analyzed by RNA Seq. Common and distinct hits were detected between the LPS and PFF-treated groups. (C) Bubble plot showing GO:CC term enrichment in the shared hits from LPS and PFFs-treated primary microglia highlights enrichment for endolysosomal processes. (D) Unsupervised hierarchical clustering shows that the treated groups cluster together suggesting common transcriptomic profiles. (E) Schematic of the in vivo experiment where LPS striatal injections were administered to WT and LRRK2 KO mice, followed by microglia isolation and single-cell RNA Seq. (F) UMAP plot showing separation of the retrieved microglia in distinct groups of activation states. (G) Microglia from LPS-injected animals spanned the activation states while PBS-injected animals gave predominantly resting microglia. (H) Lysosomal and endocytic mechanisms as well as cytoskeletal pathways are enriched in the cumulative data. (I) Heatmap showing clustering of microglia in distinct activation states highlighting increase in lysosomal and iron-related gene expression by inflammation.

We next asked whether the same regulation could be confirmed *in vivo*. Either wild type or Lrrk2 knockout mice were given a single intrastriatal injection of LPS or vehicle control and three days later adult microglia were isolated using CD11b microbeads (Fig 5E). Between 3000 and 5000 cells were recovered from each of the four groups and analyzed using single cell RNA-Seq (scRNA-Seq). We identified five distinct clusters of cells based on transcriptome similarity that represent resting microglia and four activation states (Fig 5F), with PBS-injected animals exhibiting predominantly resting microglia whereas the LPS-injected animals spanned the range of activation states (Fig 5G). Notably, we did not see strong differences between genotypes other than the KO animals tended to have fewer activated microglia, suggesting that Lrrk2 influences how microglia respond to proinflammatory stimuli (29). Gene ontology showed enrichment for lysosomal and endocytic membrane processes (Fig 5H). These analyses identified genes involved in iron storage and uptake (FTH1), vesicular transport (SNAP23), acidification of vesicles (ATP6V1G1) and the late endosomal pathway (Rab7) (Fig 5I). These data demonstrate that processes of vesicular trafficking and iron-homeostasis are modulated in a synchronized manner by inflammation. Given the importance of iron uptake in microglia activation, we hypothesized that LRRK2 may play a role in trafficking of TfR and that PD-linked mutations may affect iron uptake and accumulation in inflammatory conditions. Thus, we next sought to examine the effect of LRRK2 mutations on Tf recycling and iron uptake in proinflammatory conditions.

### G2019S LRRK2 induces transferrin mislocalization and association with lysosomes in iPSC-derived microglia

To examine whether endogenous LRRK2 mutations might influence Tf-mediated iron uptake as modulated by neuroinflammation, iPSC-derived microglia from G2019S carriers were examined in resting and proinflammatory conditions and Tf localization was assessed by super-resolution microscopy. Endogenous Tf showed partial colocalization with lysosomes in control conditions that was significantly decreased following LPS treatment, reflecting induction of rapid uptake and recycling close to the plasma membrane, in wild type cells. In contrast, the G2019S LRRK2 cells retained the association of Tf with lysosomes following LPS treatment (Fig 6A, 6B). After 3D rendering (Fig 6A, right-hand panels; suppl. video 1-4 for Imaris processing), Tf vesicle size and proximity to the nucleus were measured (Fig 6C, 6D). The G2019S LRRK2 cells showed significantly larger Tf vesicles that were sequestered closer to the nucleus compared to WT cells (Fig 6C, 6D). Additionally, in resting conditions, around 50-60% of Tf vesicles were concentrated near the juxtanuclear region, positioned within 1.5μm from the nuclear membrane, in both WT and G2019 LRRK2 cells. Upon activation by LPS, WT LRRK2 cells exhibited a more dispersed Tf localization throughout the cytoplasm with a lower fraction of vesicles close to the perinuclear recycling compartment (∼40% within 1.5μm) while in contrast, G2019S LRRK2 cells showed an increase in proximity to the nucleus (Fig 5E, 5F). These data suggest that in G2019S LRRK2 cells, Tf was retained in the perinuclear recycling compartment region associated with lysosomes, under proinflammatory conditions. A model whereby endogenous LRRK2 mutations may redirect Tf receptors to lysosomes under inflammatory conditions in microglia would be predicted to cause iron accumulation. We next tested whether we could support this hypothesis *in vivo* using the G2019S LRRK2 knock-in mouse model that expresses this mutation in the appropriate endogenous context.

**Figure 6.**
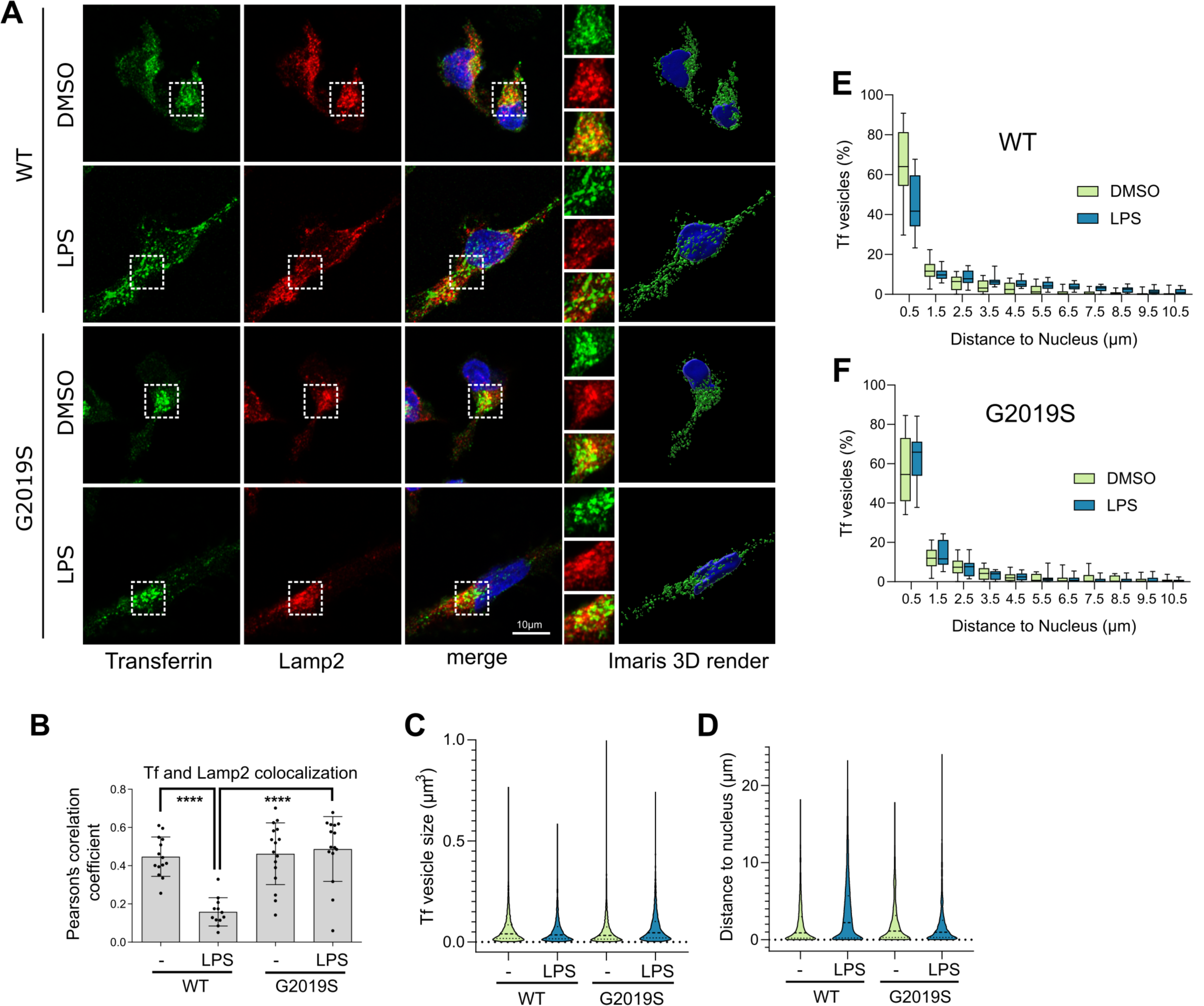
G2019S LRRK2 modulates transferrin recycling in iPSC-derived human microglia. (A) iPSC-derived human microglia from WT or G2019S LRRK2 carriers were treated with LPS and the localization of transferrin (Tf) and Lamp2 were analyzed by super-resolution microscopy and surface rendering in Imaris (A). Partial co-localization between Tf and Lamp2 was observed in control that was significantly decreased with LPS treatment in WT cells but not in G2019S LRRK2 cells that retained lysosomal association of Tf (B) (N=16, one-way ANOVA Tukey’s post hoc, ****P<0.0001, F(3,53)=16.16). G2019S LRRK2 iPSC microglia exhibited larger Tf vesicles compared to WT while LPS treatment induced a decrease in average vesicle size in both cohorts (C) (minimum 3000 vesicles were counted from 16 cells per group, two-way ANOVA, genotype: P<0.0001, F(1,17753)=16.38, treatment: P<0.0001, F(1,17753)=20.57). LPS treatment induced an increase in the average distance of Tf vesicles from the nucleus in WT cells but that was not significant in G2019S LRRK2 cells (D) (minimum 3000 vesicles were counted from 16 cells per group, two-way ANOVA, genotype: P<0.0001, F(1,14735)=104.9, treatment: P<0.0001, F (1, 14735) = 105.0). The frequency distributions of Tf vesicle proximity to the nucleus were plotted in E and F. The percentage of Tf vesicles proximal to the nucleus was significantly decreased with LPS treatment in WT but not in G2019S LRRK2 cells (E, F) (E: bin at 0.5μm, two-tailed student’s t-test; Matt-Whitney post-hoc; **P=0.0022; F: bin at 0.5μm, two-tailed student’s t-test; Matt-Whitney post-hoc; NS).

### G2019S LRRK2 induces iron and ferritin accumulation in inflammatory microglia *in vivo*

To examine the effect of LRRK2 mutations on iron accumulation in inflammation, we again used intrastriatal LPS injections on age-matched WT, Lrkk2 KO and G2019S knock-in mice. Perls staining was used to visualize iron deposition in sections that spanned the striatum and substantia nigra 72 hours after injection (Fig 7A). The G2019S LRRK2 knock-in mice showed a marked increase in iron deposition in the striatum compared to WT and Lrrk2 KO mice as analyzed by densitometry (Fig 7B, 7C). Higher magnification images revealed that the Perls stained cells had a microglial morphology, in line with out *in vitro* data that suggest dyshomeostasis of iron regulation mechanisms in inflammatory microglia (Fig 7D). Additional sections from the same animals were stained for FTH1 and Tf. Accumulation of ferritin was observed in G2019S LRRK2 mice compared to WT and LRRK2 KO mice, with patterns similar to that seen with iron deposition (Fig 8A, 8B) while Tf was not significantly altered in the knock-in model (Fig 8C). To characterize further the type of cells that accumulate ferritin, sections were co-stained for Iba1 and analyzed by imaging (Fig 8D). Iba1-positive microglia accumulated ferritin in both WT and G2019S LRRK2 LPS-treated cohorts. About 50% of ferritin-positive cells were Iba1-positive in both groups (Fig 8E). These data suggest that in G2019S LRRK2 knock-in mice, striatal microglia accumulate higher levels of ferritin-bound iron upon inflammation by LPS compared to WT LRRK2 mice.

**Figure 7.**
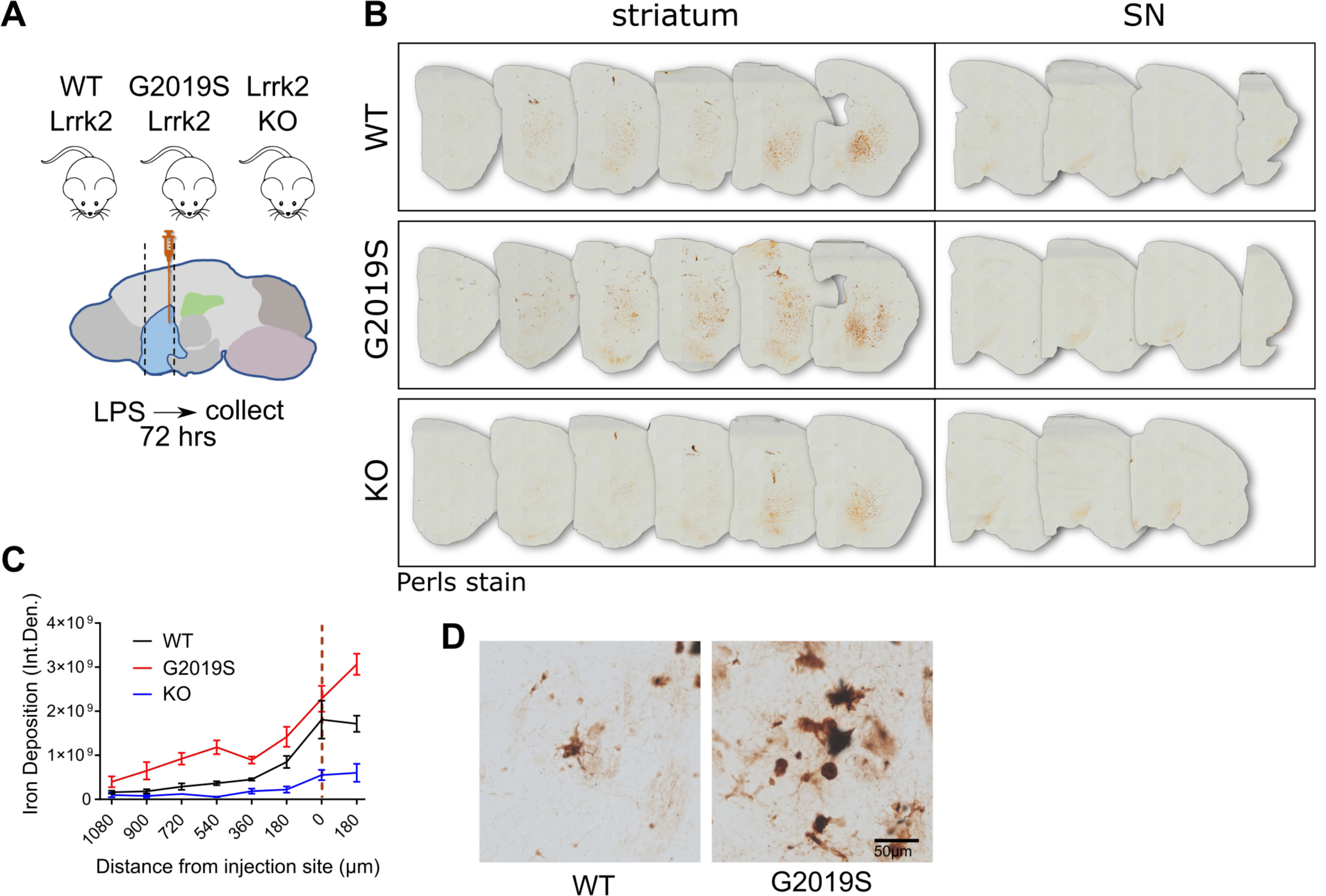
G2019S LRRK2 induces iron accumulation in inflammatory microglia *in vivo*. (A) Schematic of experimental design: WT, G2019S knock-in and Lrrk2 KO mice were administered intrastriatal injections of LPS and 72 hours later brains were collected and stained by Perls stain. (B, C) G2019S LRRK2 knock-in mice exhibited significantly higher iron deposition in the striatum proximal to the injection site compared to WT and Lrrk2 KO, while minimal signal was detected in the substantia nigra in all groups (two-way ANOVA, Tukey’s post-hoc, Genotype: ***P=0.0007, F (2, 7) = 24.70, Distance from injection site: ****P<0.0001, F (7, 49) = 32.28). High-magnification images revealed iron deposition in inflammatory microglia (D). [SEM bars are shown]

**Figure 8.**
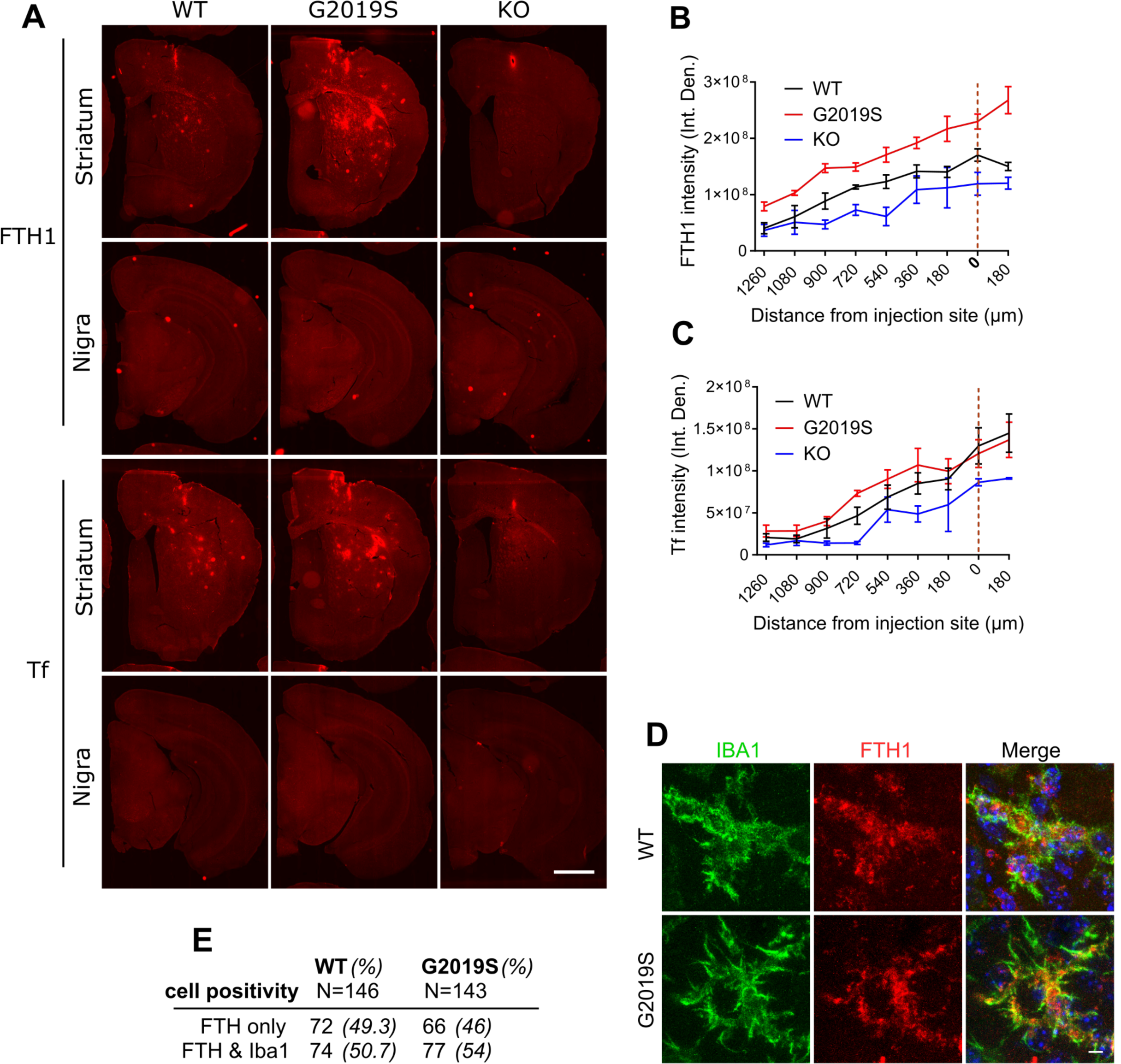
Inflammation induces ferritin accumulation in microglia in G2019S LRRK2 knock-in mice. Ferritin heavy chain (FTH) and transferrin (Tf) were stained and visualized in collected brains, 72 hours after post intrastriatal injections of LPS. G2019S LRRK2 mice exhibited higher levels of FTH across the striatum compared to WT and Lrrk2 KO mice while Tf was not altered significantly in the knock-in (A, B, C) (B: two-way ANOVA, Tukey’s post-hoc, Genotype: **P=0.0017, F (2, 7) = 18.27, Distance from injection site: ****P<0.0001, F (8, 56) = 50.25; C: two-way ANOVA, Tukey’s post-hoc, Genotype: P=0.1040, F (2, 7) = 3.182, Distance from injection site: ****P<0.0001, F (8, 56) = 29.17). Microglia are positive for FTH in both WT and G2019S LRRK2 cohorts (D). Similar percentages of FTH-positive microglia were observed between the WT and G2019S LRRK2 groups (E). [SEM bars are shown].

## Discussion

In this study, we show that PD-linked LRRK2 mutations have a convergent phenotype of Rab8a mislocalization away from the ERC and recruitment to damaged lysosomes that is distinct from wild type protein. This relocalization is associated with dysregulation of Rab8a-mediated transferrin recycling both in heterologous cell lines and in human iPSC-derived microglia from PD patients with LRRK2 mutations following inflammatory stimulus (Fig 9 for model schematic). Furthermore, we show that in G2019S LRRK2 knock-in mice, LPS-induced inflammation in the striatum results in higher iron accumulation in microglia and increased ferritin staining *in vivo*. These results suggest that LRRK2 plays a role in iron homeostasis response in neuroinflammation, driven, at least in part, by altered endolysosomal functions.

**Figure 9.**
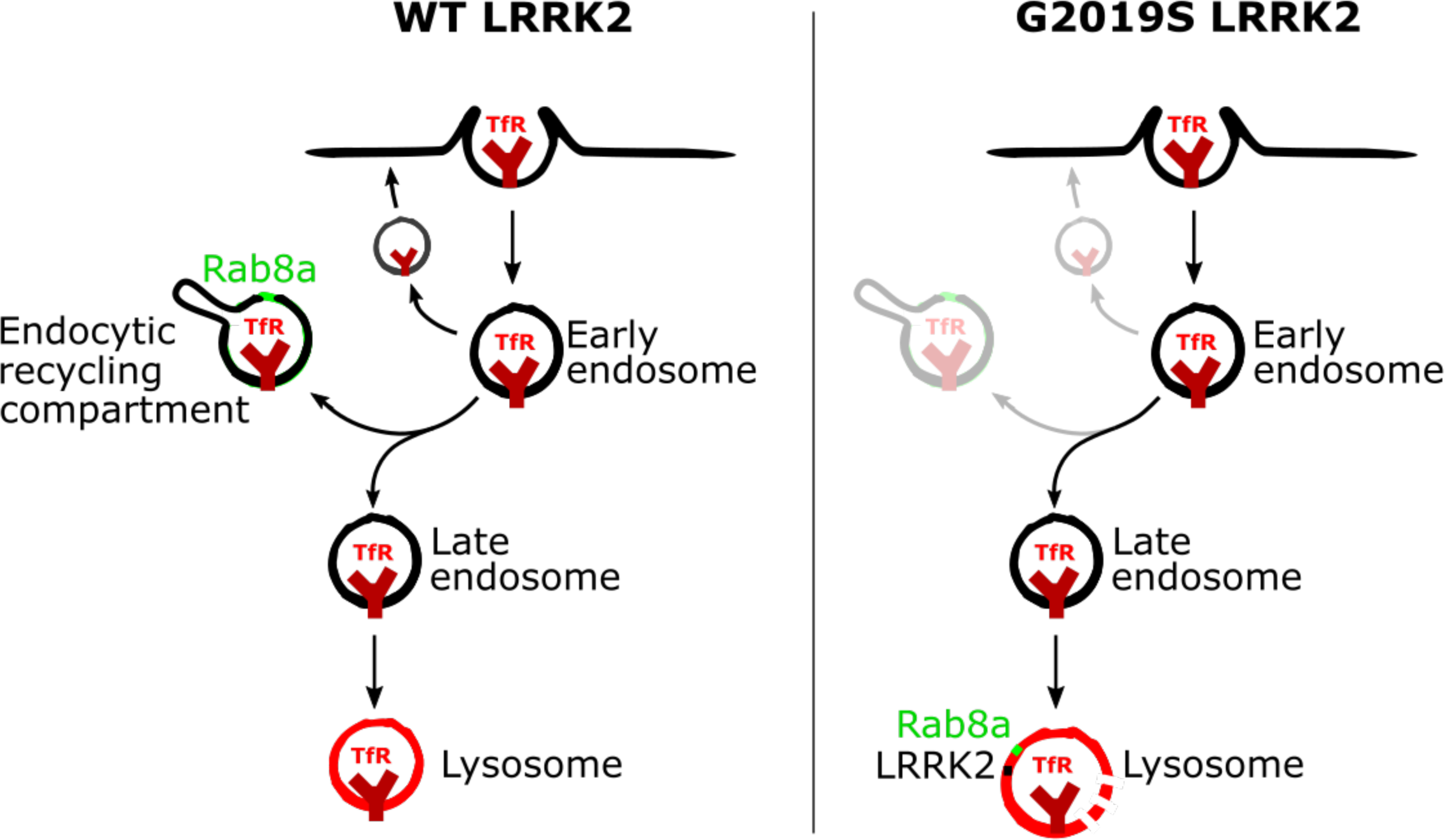
A model of the recruitment of Rab8a to lysosomes and dysregulation of transferrin recycling by mutant LRRK2. Following clathrin-mediated endocytosis, TfR can undergo rapid recycling close to the membrane, recycling via the ERC or targeting to lysosomes. Rab8a mediates recycling of TfR via the ERC and LRRK2 mutations that hyperphosphorylate Rab8a induce the sequestration of Rab8a and Tf to damaged lysosomes.

Transferrin-mediated iron uptake is the main route of iron delivery to most cell types. Extracellular transferrin binds ferric iron (Fe3+) and is internalized by TfR through clathrin-mediated endocytosis. TfR is trafficked through early endosomes and can either be rapidly recycled back to the membrane or associate with the ERC in a slower recycling step. In the acidic lumen of endolysosomes, ferric iron (Fe3+) is reduced to ferrous iron (Fe2+), which mediates its release from Tf and the export to the cytosol through DMT1, where iron is subsequently delivered to different subcellular components. Lysosomal pH regulates iron release and impaired lysosomal acidification has been reported to trigger iron deficiency and inflammation *in vivo* as well as mitochondrial defects (30,31). As LRRK2 mutations have been reported to alter the autophagic lysosomal pathway in carriers (32,33), as well as to perturb lysosomal acidification in knock-in mouse models (34), it is plausible that LRRK2 mutations affect Tf-mediated iron uptake by modulating lysosomal acidification. In our experiments we observe sequestration of TfR and accumulation to lysosomes driven by mutant LRRK2, while no significant difference was found on total TfR levels in these cells (data not shown) or in G2019S knock-in mice that exhibited iron deposition and ferritin upregulation following LPS administration. Consistent with these observations, transferrin receptor levels are not being found to be increased in PD or in mouse models of the disease (35,36) suggesting that the regulation of transferrin recycling through the endolysosomal system rather than overall levels contribute to iron dyshomeostasis. LRRK2 deficiency also reportedly impairs recycling of TfR and is required to maintain the lysosomal degradative capacity of endocytic and autophagic cargo (37). The LRRK2 G2019S mutation can affect Rab8a-mediated receptor recycling and endolysosomal transport (26). Lastly, we have recently shown that LRRK2 mutations can impair clathrin-mediated endocytosis, which governs internalization of different receptors including TfR (38). Therefore, the effect of LRRK2 mutations on TfR recycling may be driven by a dysregulation of vesicular trafficking through different Rab GTPases.

Iron deposition in the brain is a feature of PD and other neurodegenerative diseases. Imaging and biochemical methods have confirmed iron accumulation in the SN of PD patients correlating with severity of motor symptoms (39–42). Iron deposition can be detected by transcranial sonography in post-mortem brains from PD patients that correlates with increased ferritin levels and loss of neuromelanin content as assessed biochemically (43). Studies on animal models have also supported a role of iron in nigral neurodegeneration in PD. In a model of acute MPTP intoxication in mice, dopaminergic cell loss correlated with iron accumulation and increase in lipoperoxidation coinciding with upregulation of the divalent metal transporter DMT1 (44). Furthermore, iron chelators can rescue dopaminergic neuron loss and behavioral effects caused by intracerebroventricular administration of 6-hydroxydopamine in rats (45). In the case of LRRK2, higher nigral iron deposition has been reported in LRRK2 mutation carriers compared to idiopathic patients, while also the same trend has been highlighted in Parkin mutation carriers (46). Pink1 and Parkin have been linked to degradation of mitochondrial iron importers (47,48) while iron overload can induce a Pink1/Parkin-mediated mitophagic response (49). These data identify convergent pathways that regulate iron homeostasis and that may be involved in PD pathogenesis in patients.

We show that in G2019S LRRK2 knock-in mice, LPS-induced inflammation in the striatum results in higher iron accumulation in microglia compared to WT mice. Studies have reported a cooperative effect of neuroinflammation and iron accumulation. Microglial activation by LPS induces secretion of IL-1β and TNF-α that in turn activate iron regulatory proteins in dopaminergic neurons inducing iron overload and neurotoxicity (50). In these experiments the iron status of microglia exacerbated proinflammatory cytokine release and neuronal degeneration. Furthermore, in mixed-culture models, microglia play a pivotal role in iron-elicited dopaminergic neurotoxicity via increase in cytokine production (51). The iron storage protein ferritin is reportedly decreased in the SN of PD patients compared to controls (39). Increased iron deposition together with lower ferritin levels could indicate increase in intracellular labile iron pools that constitute chelatable redox-active iron damaging to cells. Transgenic mice over-expressing human ferritin protein in dopaminergic SN neurons do not exhibit increases in reactive oxygen species or SN neuron loss following systemic administration of MPTP (52). In G2019S LRRK2 knock-in mice we observe an increase in ferritin that may represent a compensatory event to limit neurotoxicity, as a result of dysregulated Tf recycling and iron accumulation.

Iron overload in the brain can activate glial cells and promote the release of inflammatory and neurotrophic factors that control iron homeostasis in dopaminergic neurons. Increased iron in neurons can be toxic through production of hydroxyl radicals via Fenton chemistry, responsible for oxidation of lipids, proteins, and DNA. LRRK2 is modulated by oxidative stress in cells and PD-linked mutations compromise mitochondrial integrity (53–55). Recent studies have demonstrated direct delivery of iron from Tf-endosomes to mitochondria via “kiss-and-run” events while highlighting how these events sustain mitochondrial biogenesis (56). Dysregulation of iron storage or uptake by LRRK2 in microglia may in turn affect cytokine production and neuronal survival, as well as have a feedback effect on LRRK2 activity in the brain.

We have not yet established whether the observed effects of LRRK2 on iron homeostasis and transferrin recycling are damaging in the context of disease. This is due to the limitations of the available knock-in animal models that they do not present with baseline neurodegeneration. As a consequence of this, it remains uncertain whether iron modulation is a promising therapeutic avenue in the context of LRRK2 mutation carriers. Iron chelators have proven promising in neurodegeneration with brain iron accumulation disorders (57), but iron chelation therapy did not improve motor-UPDRS scores and quality of life significantly (58). In recent years, LRRK2 has been nominated to play roles in a number of cellular processes ranging from inflammation, autophagy and endolysosomal pathways to mitochondrial homeostasis, processes that may also be cell-type specific. Given the role that iron dyshomeostasis seems to play in basal ganglia diseases, a combination therapy of iron chelation along with inhibition of LRRK2 could present a viable strategy.

Our data suggest an effect of LRRK2 mutations on Rab8a function, driven by increased kinase activity, that may drive a dysfunction of iron-uptake mechanisms in response to inflammatory stimuli in resident microglia. Deciphering the protein trafficking pathways around LRRK2 will help us understand the mechanistic underpinnings of neurodegeneration and the biological implications of blocking LRRK2 kinase in the clinic, while highlighting signaling avenues that can be targeted as therapeutic means.

## Materials and Methods

### Cell Culture, Treatments, and Constructs

HEK293FT cells (Thermo Scientific) were cultured in DMEM supplemented with 10% FBS and maintained at 37°C, 5% CO2. HEK293T cell lines stably expressing different GFP LRRK2 variants were grown and cultured as described previously (59). HEK293FT cells were transfected using Lipofectamine 2000 using standard procedures. For siRNAs transfection, cells were transfected with the SMARTpool ON-TARGETplus or scrambled siRNA control (Dharmacon) using the DharmaFECT reagent, according to the manufacturer’s instructions. The 3×FLAG-tagged construct of LRRK2 in pCHMWS plasmid was a gift from Dr. J. M. Taymans (KU Leuven, Belgium).

### Structure modelling

Heterodimeric complexes (Rab8a and Rabin8) and (Rab8a and Mical-L1) were modeled in PyMol (PDB: 4LHY and 5SZH respectively) (60–62). Insertion of the phosphate at T72 was modeled and resulting distances were measured using the tools available (PyMOL version 2.0).

### Animal procedures

C57BL/6J mice were housed in standardized conditions at 2-5 animals per cage and with ad libitum access to food and water on a 12h light-dark cycle. All procedures with animals followed the guidelines approved by the Institutional Animal Care and Use Committee of National Institute on Aging, NIH.

### Stereotaxic surgery

One-year old WT, LRRK2 KO or LRRK2 G2019S mice were kept under anesthesia using 1-2% isoflurane. Mice were placed into a stereotaxic frame, an incision was made above the midline and the skull was exposed using cotton tips. At anteroposterior +0.2 mm, mediolateral +/-2.0 mm from bregma (bilateral injection), a hole was drilled into the skull. A pulled glass capillary (blunt) attached to a 5μl Hamilton glass syringe was used for injecting either 1 μl of either PBS or 5mg/ml LPS solution (5µg) per hemisphere. The capillary was lowered to dorsoventral −3.2 mm from bregma into the dorsal striatum. The solution was delivered at a rate of 0.1 μl per 10sec. After the injection, the capillary was held in place for 2 min, retracted 0.1 μm and another 1 min was waited before it was slowly withdrawn from the brain. The head wound was closed using surgical staples. Ketoprofen solution at 5mg/kg was administered subcutaneously as analgesic treatment for the following 3 days.

### Histology

Animals were sacrificed 3 days after surgery. Mice were deeply anesthetized using an i.p. injection of 200ul of 10% ketamine. The thoracic cavity was opened to expose the heart. The whole body was perfused with 10 ml of 0.9 % NaCl (2min). Brains were removed, the left hemisphere was used for WB analysis while the right hemisphere was fixed in 4 % PFA for 48 h. After 2 days, fixed hemispheres were transferred to 30 % sucrose solution for cryoprotection. The brains were cut into 30 μm thick coronal sections - 6 series - and stored in antifreeze solution (0.5 M phosphate buffer, 30 % glycerol, 30 % ethylene glycol) at −20°C until further processed.

### Immunohistochemistry

Sections were washed with PBS and incubated for 30min in blocking buffer (10% [Normal Donkey Serum] NDS, 1% BSA, 0.3% Triton in PBS). Afterwards, primary antibodies rabbit ant-FTH1 (D1D4; 4393S, Cell Signalling) and goat anti Iba1 (abcam, ab5076) were used at 1:500 and incubated overnight at 4°C in 1%NDS, 1% BSA, 0.3% Triton in PBS. Next day, sections were washed 3x for 10min each with PBS and incubated with Alexa Fluorophore (568 or 647) conjugated secondary antibodies for 1h at room temperature. After 3 washes with PBS, sections were mounted on glass slides, coverslipped using Prolong Gold Antifade mounting media (Invitrogen) and imaged using a Zeiss LSM 880 confocal microscope equipped with Plan-Apochromat 63X/1.4 numerical aperture oil-objective (Carl Zeiss AG). Sections were further imaged using an Olympus VS120 (Olympus, Center Valley, PA) slide scanner microscope.

### Perls Blue staining

Sections were washed in ddH2O 3x for 10min each. Afterwards, sections were incubated in a 1:1 mix of 4% potassium ferrocyanide (Sigma-Aldrich P3289-100G) and 4% HCl. After 30min, brain slices were washed 3x for 10min with PBS and quenched using 10% Methanol + 3% H2O2 diluted in PBS for 1h. Sections were then rinsed again using PBS and incubated in 3,3′-Diaminobenzidine and H2O2 according to instructions (SIGMAFAST D4418-50SET, Sigma-Aldrich). Then, slices were washed with PBS 3x for 10min, mounted on SuperFrost Plus slides (Fisher Scientific), dried overnight and dehydrated using 70% Ethanol, 95% Ethanol, 100% Ethanol followed by Xylene and DPX mounting media (Sigma, 06522). and analyzed on ImageJ for signal intensity of selected ROIs.

### ICPMS

Total iron concentrations in the samples were measured by ICP-MS (Agilent model 7900). For each sample, 200 µL of concentrated trace-metal-grade nitric acid (Fisher) was added to 25 µL or 100 µL of sample taken in a 15 mL Falcon tube. Tubes were sealed with electrical tape to prevent evaporation, taken inside a 1L glass beaker, and then placed at 90 0 C oven. After overnight digestion, each sample was diluted to a total volume of 4 mL with deionized water, and then analyzed by ICP-MS.

### iPSC differentiation

Induced pluripotent stem cell (iPSC) lines were derived from reprogrammed peripheral mononuclear blood cells (PBMCs) collected from participants of the Parkinson’s Progression Marker Initiative (PPMI). Differentiation of iPSC to microglia was accomplished via a hematopoietic stem cell intermediate stage according to published protocols (63,64). Mature iPSC-derived microglia were characterized by western blot and qPCR for expression of Iba1 protein and AIF1, TMEM119 and P2RY12 mRNA, respectively. (data not shown). The lines used were PPMI 3448 (WT) and PPMI 51782 (G2019S LRRK2 carrier). LPS activation was done in differentiation media at 100ng/ml for 16hr prior to fixation and staining. Activation by LPS treatment was verified by WB of pNFKb, p38 and imaging of Iba1 staining (Fig S4).

### Co-immunoprecipitation

HEK-293T were transfected with 3×FLAG-Rab8a variants and 3xFLAG-GUS (Fig 3D) or 3xFLAG LRRK2 variants and GFP-Rab8a (Fig 3E), using Lipofectamine 2000 as per the manufacturer’s instructions. After 24 h, cells were lysed in buffer containing 20 mM Tris/HCl (pH 7.4), 137 mM NaCl, 3 mM KCl, 10% (v/v) glycerol, 1 mM EDTA, and 0.3% Triton X-100 supplemented with protease inhibitors and phosphatase inhibitors (Roche Applied Science).

Lysates were centrifuged at 21,000 × g, 4 °C for 10 min, and the supernatants were analyzed for protein concentration (Pierce). 5 μg of total protein from each supernatant was analyzed by SDS-PAGE for expression of the proteins in question. For FLAG immunoprecipitations, 3 mg of each sample was precleared with EZview protein G beads (Sigma-Aldrich) for 0.5 h at 4 °C, and subsequently FLAG M2 beads were incubated with the lysates for 2 h at 4 °C to IP target constructs. FLAG-tagged proteins were eluted in 1x kinase buffer (Cell Signaling), containing 150 mM NaCl, 0.02% Triton and 150 ng/µl of 3xflag peptide (Sigma-Aldrich) by shaking for 30 minutes at 4°C. For GFP IPs, Chromotek-GFP-Trap-agarose resin (Allele Biotech) was incubated with lysates for 2 h at 4 °C. The beads were washed four times with buffer containing 20 mM Tris/HCl (pH 7.4), 137 mM NaCl, 3 mM KCl, and 0.1% Triton X-100. The washed beads were boiled for 6 min in 4× NuPAGE loading buffer (Invitrogen) supplemented with 1.4 M β-mercaptoethanol and analyzed by Western blot.

### Western Blot

Standard Western blot protocols were used with the following antibodies: anti-LRRK2 (ab133474; Abcam), anti-Rab8a (6975, Cell Signaling), anti-MICAL-L1 (H00085377, Abnova), GDI1/2 (716300, Thermo Scientific), Rabin8 (12321-1-AP, Protein Tech), FLAG (F1804, Sigma-Aldrich).

### RNA isolation and RNA-seq

For figure 5A-D, RNA was isolated from primary microglia and analyzed by bulkRNA sequencing following methods published in (28). For figure 5E-I, mice were administered striatal LPS injections and sacrificed 3 days later. Collected brain hemispheres were dissociated using the Adult Brain Dissociation kit (Miltenyi Biotec) and brain microglia cells were isolated using CD11b microbeads and MACS sorting columns (Miltenyi Biotec). Single cell suspensions were subjected to scRNA-seq at the National Cancer Institute single cell analysis Core facility, Bethesda, MD and followed the 10xGenomics pipeline. Data were analyzed using the R package Seurat (version 3, PMIDS: 26000488; PMID: 29608179) as described in (28).

### Immunocytochemistry

HEK293fT cells were seeded at 0.1 × 10^6^ cells/well on 12mm coverslips precoated with poly-D-lysine (Millicell EZ slide, Millipore) and cultured as described before above. Cells were fixed in 4% (w/v) formaldehyde/PBS, blocked in 5% (v/v) FBS in PBS, and incubated with primary antibodies in 1% (v/v) FBS/PBS for 3 hrs. Following three washes in PBS, the cells were incubated for 1 h with secondary antibodies (Alexa Fluor 488, 568, 647-conjugated; ThermoFisher). After three PBS washes, the coverslips were mounted and the cells were analyzed by confocal microscopy (Zeiss LSM 880 and Airyscan super-resolution microscopy). For high-content imaging, where Rab8a translocation was quantified, cells were plated in 96-well plates, precoated with matrigel, transfected with LRRK2 mutants and processed following 24 hrs of expression. Cells were treated with DMSO or 1µM MLi-2 for 1hr prior to fixation and staining with the corresponding antibodies. Cells were visualized with the ThermoFisher Cellomics ArrayScan using the HCS Studio platform and the SpotDetector v4 bioassay protocol. A minimum of 800 cells were imaged per well from a total of 6 wells per construct and condition and Rab8a localization was analyzed.

### Imaris (Bitplane) analysis

Following super-resolution microscopy, the Imaris (Bitplane) platform was used to analyze the localization of TfR and Tf in different experiments. In figure 4A, TfR staining was rendered to spots throughout z planes in the 3D volume, and the distance between them was counted. In figure 6, super-resolution z-stack images of iPSC-derived microglia were used to render Tf and nuclear staining to surfaces and measure the distance of Tf vesicles from the nucleus edge (3D rendering process outlined in supplementary videos 1-4).

### Transferrin Recycling Assay

Cells that were plated and transfected on 12mm poly-D-lysine coated coverslips were processed following 24 hrs of expression. Initially, cells were incubated in pre-warmed DMEM for 45 mins. Alexa 568 fluorophore-conjugated transferrin (T23365, ThermoFisher) was added to the media at 20µg/ml final concentration and cells were incubated for 30 mins. Following one quick wash in prewarmed media, cells were incubated with prewarmed DMEM, 10% (v/v) FBS media for 10 mins and fixed and stained as detailed above. For the uptake assay, cells were plated in 96-well plated (15,000 cells/well), transiently transfected and processed the next day. Cells were transferred to the ThermoFisher Cellomics ArrayScan and incubated in DMEM for 45 mins. Equal volume of DMEM media was added to wells containing 40µg/ml of Alexa 568 fluorophore-conjugated transferrin, to achieve 20µg/ml final transferrin concentration, and cells were imaged at 0, 2, 4, 6, 8, 10, 15 and 20 mins. A minimum of 800 cells were imaged per well, in a total of 6 wells per condition and intracellular transferrin levels were quantified.

### Statistical Analysis

Statistical tests used are noted in figure legends of representative graphs. Briefly, one-way ANOVA or two-way ANOVA with Tukey’s post hoc-test, and two-tailed unpaired student t-test with Mann Whitney U post-hoc test were used. All statistical analyses were performed using GraphPad Prism 7 (GraphPad Software, San Diego, CA). Experiments were done in 3 independent experimental replicates unless otherwise indicated.

## Acknowledgements

We thank the NCI-CCR Single Cell Analysis Facility, Cancer Research Technology Program at the Frederick National Lab for Cancer Research for critical assistance with single cell RNA-Seq analysis. We thank Dr George R. Heaton (Icahn School of Medicine at Mount Sinai) for critical discussions on experimental procedures and data analysis.

## Supplementary Material

**Supplementary Figure S1. Validation of Rab8a antibodies for endogenous detection**. (A) The rabbit anti-Rab8A antibody by Cell Signaling (D22D8; #6975) was validated for detection of endogenous Rab8a by immunocytochemistry and western blot, in Hek293T cells transfected with NTC or Rab8a siRNA. (B) In contrast, the mouse anti-Rab8a (#ab128022) and rabbit anti-Rab8a (#ab188574) antibodies failed to validate by siRNA in cell staining and were not used for subsequent experiments.

**Supplementary Figure S2. LRRK2 genetic variants sequester endogenous Rab8a to lysosomes in a kinase-dependent manner**. Endogenous Rab8a is sequestered to Lamp2-positive lysosomes in cells expressing R1441C, Y1699C, G2019S, or I2020T LRRK2, but not the kinase-dead K1906M LRRK2 or transfection control (GUS). MLi-2 treatment rescues this phenotype.

**Supplementary Figure S3. Rab8a, TfR and LRRK2 are sequestered to damaged lysosomes**. HEK293T cells stably expressing WT LRRK2 were treated with LLOMe for 2hrs and stained. LLOMe treatment induced the sequestration of Rab8a to LRRK2 positive vesicles with partial recruitment of TfR.

**Supplementary Figure S4. Characterization of iPSC-derived microglia**. (A) Immunoblot analysis of iPSC-derived microglia following treatment with 100ng/mL LPS. (B, C) Quantification of indicated phospho-proteins normalized against total protein and loading control b-actin in treated vs. untreated cells (technical n = 3, unpaired t-test, p < 0.01). (D) Iba1 staining of DMS and LPS treated microglia from WT and G2019S cell lines.

**Supplementary Movies 1-4. Imaris 3D surface rendering of endogenous transferrin and Lamp2 staining in WT and G2019S LRRK2 iPSC-derived microglia following DMSO or LPS treatment**.

