## Supplementary figures and images for "Pathogenic mutations in LRRK2 sequester Rab8a to damaged lysosomes and regulate transferrin-mediated iron uptake in microglia"

### Fig S1

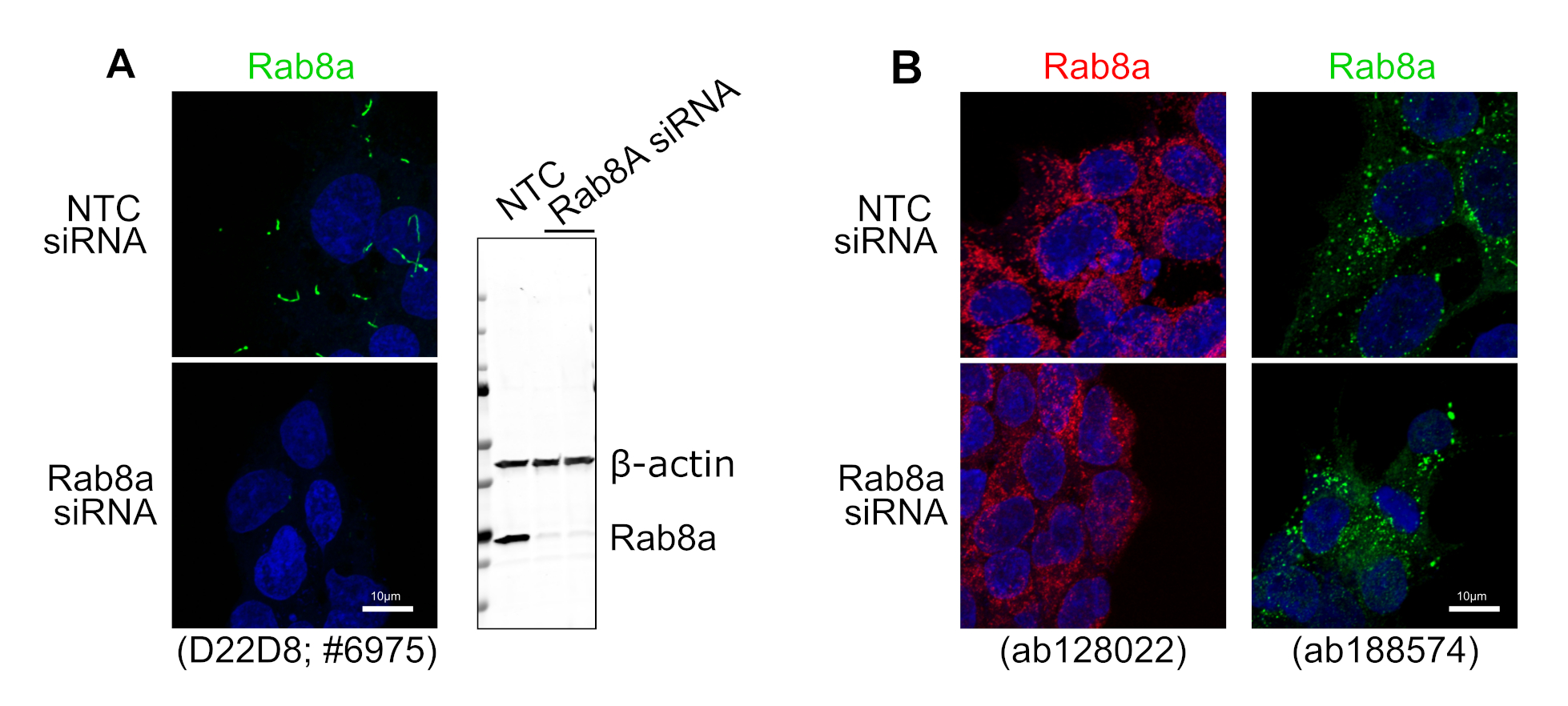

### Fig S2

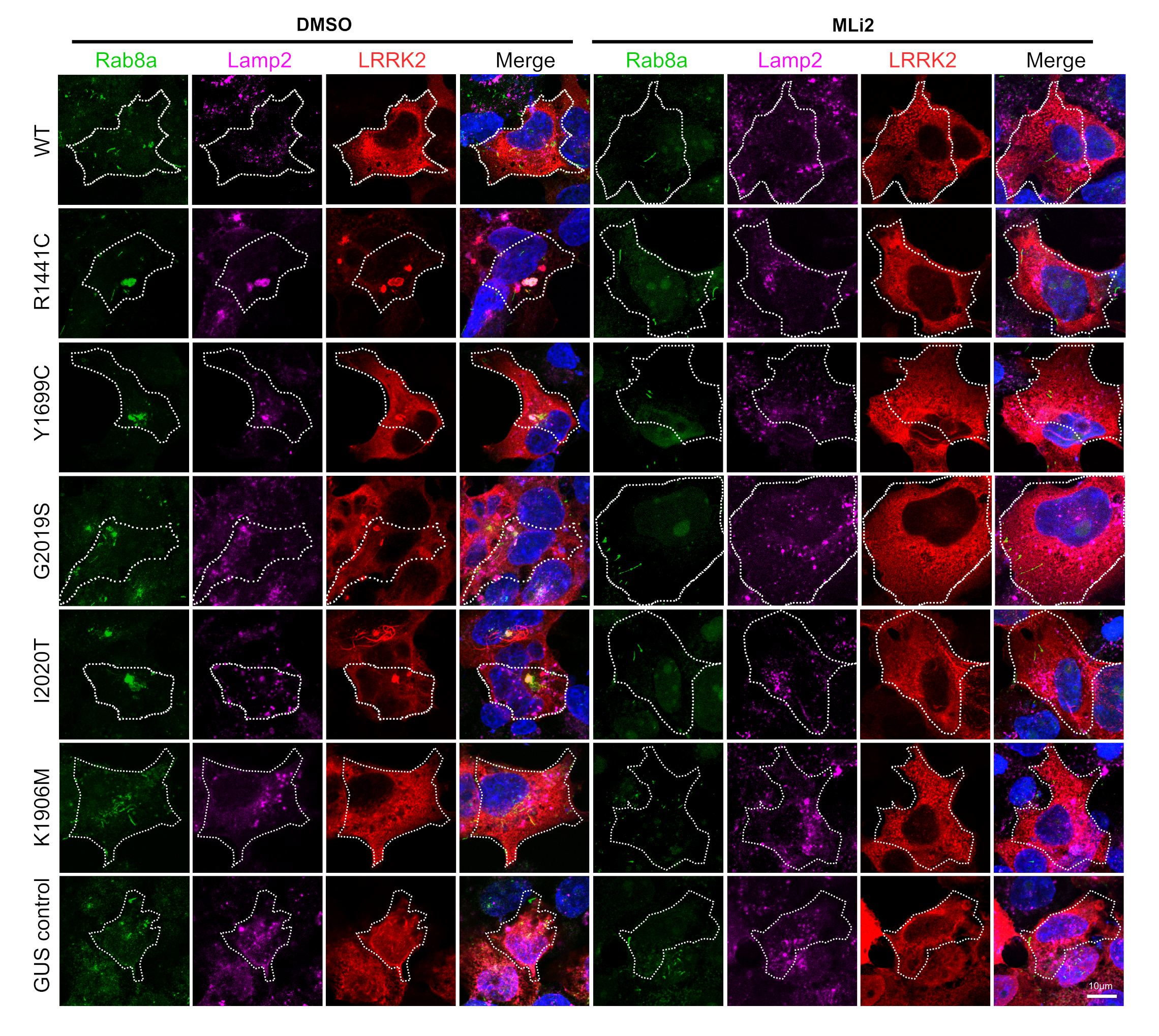

### Fig S3

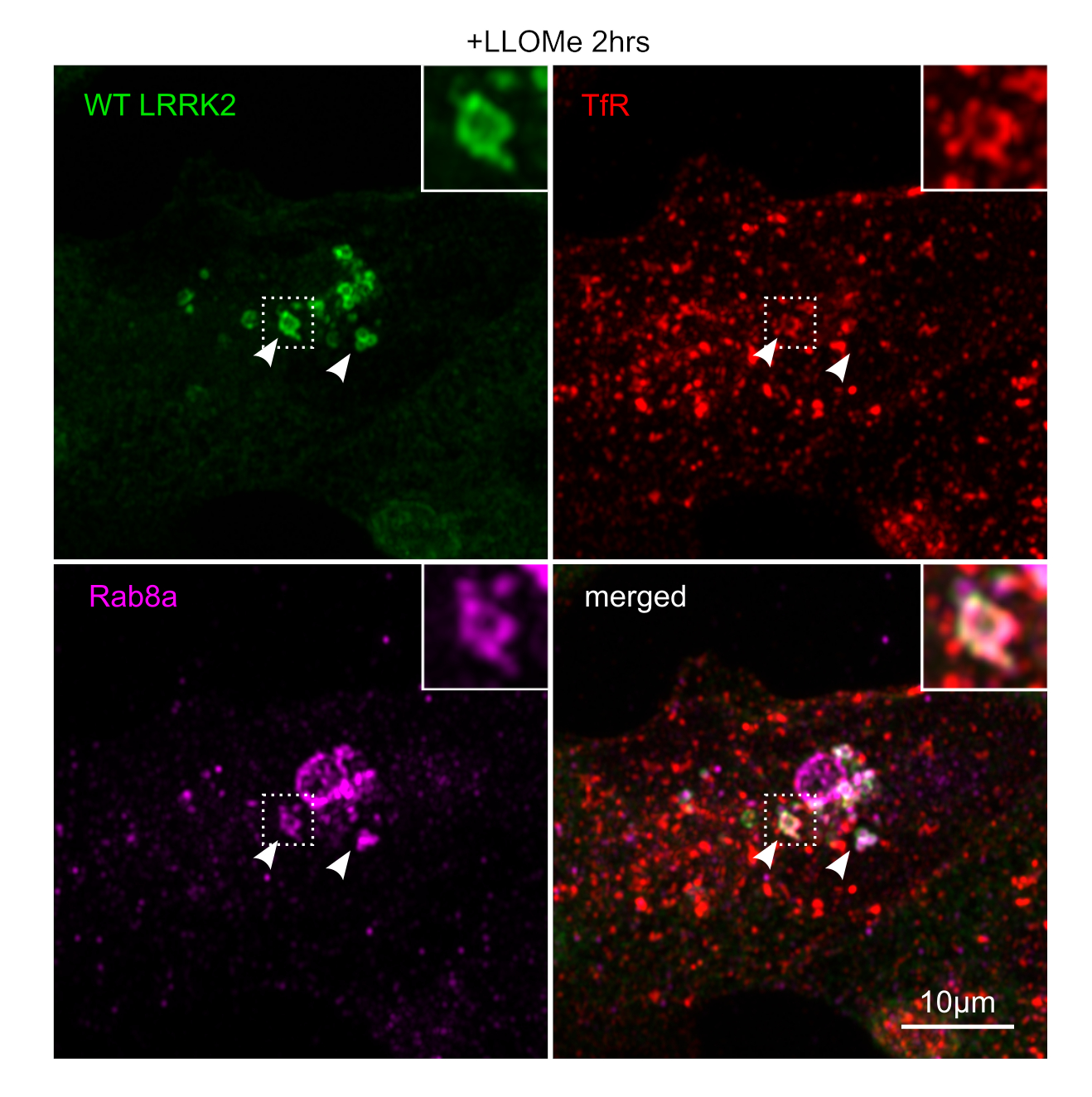

### Fig S4

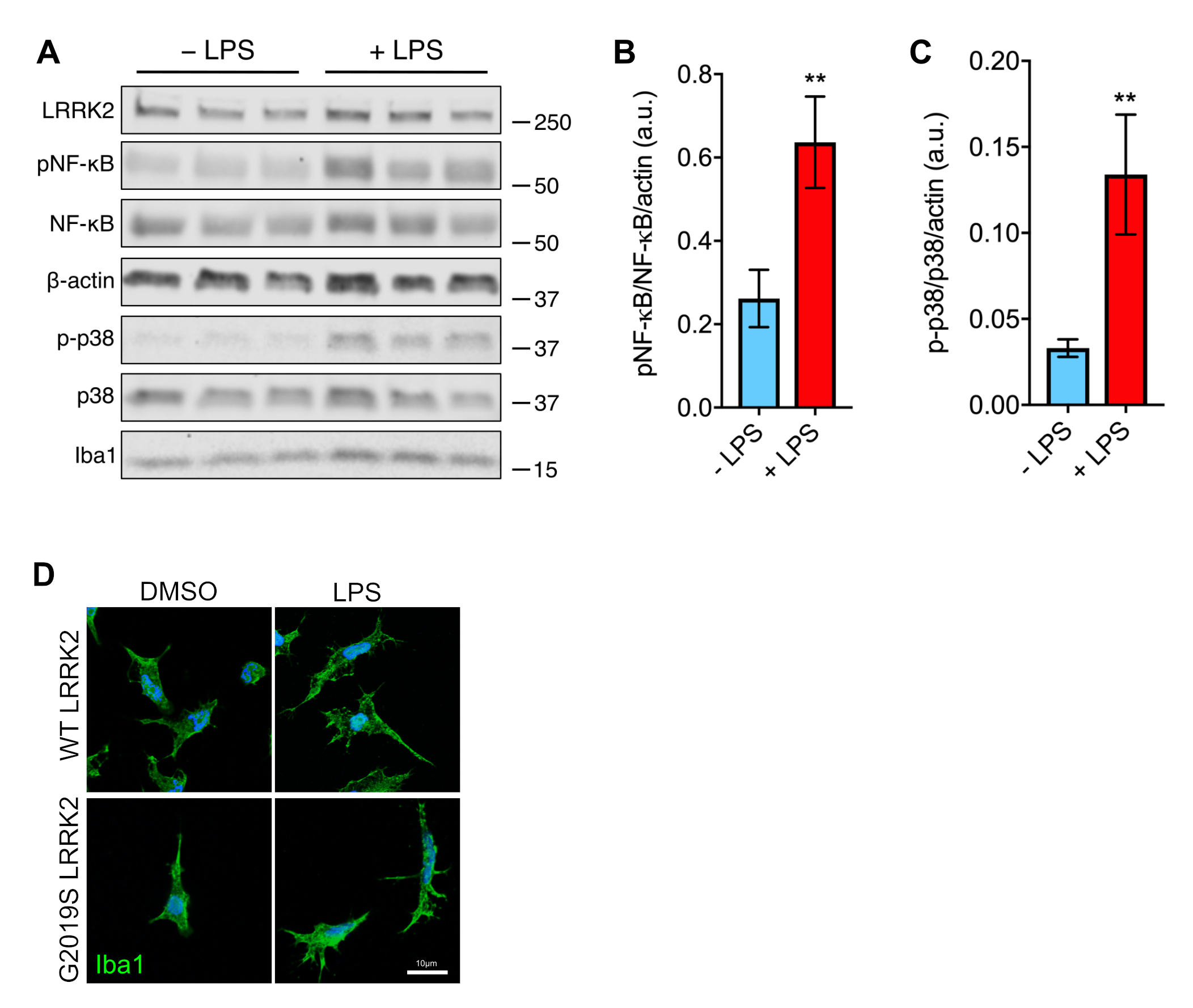
